# Construction of High-Resolution RAD-Seq Based Linkage Map, Anchoring Reference Genome, and QTL Mapping of the Sex Chromosome in the Marine Medaka *Oryzias melastigma*

**DOI:** 10.1101/695304

**Authors:** Bo-Young Lee, Min-Sub Kim, Beom-Soon Choi, Atsushi J. Nagano, Doris Wai Ting Au, Rudolf Shiu Sun Wu, Yusuke Takehana, Jae-Seong Lee

## Abstract

Medaka (*Oryzias* spp.) is an important fish species in ecotoxicology and considered as a model species due to its biological features including small body size and short generation time. Since Japanese medaka *Oryzias latipes* is a freshwater species with access to an excellent genome resources, the marine medaka *Oryzias melastigma* is also applicable for marine ecotoxicology. In genome era, a high-density genetic linkage map is a very useful resource in genomic research, providing a means for comparative genomic analysis and verification of *de novo* genome assembly. In this study, we developed a high-density genetic linkage map for *O. melastigma* using restriction-site associated DNA sequencing (RAD-seq). The genetic map consisted of 24 linkage groups with 2,481 RAD-tag markers. The total map length was 1,784 cM with an average marker space of 0.72 cM. The genetic map was integrated with the reference-assisted chromosome assembly (RACA) of *O. melastigma*, which anchored 90.7% of the assembled sequence onto the linkage map. The values of complete Benchmarking Universal Single-Copy Orthologs (BUSCO) were similar to RACA assembly but N50 (23.74 Mb; total genome length 779.4 Mb; gap 5.29%) increased to 29.99 Mb (total genome length 778.7 Mb; gap 5.2%). Using MapQTL analysis with a single nucleotide polymorphism markers, we identified a major quantitative trait locus for sex traits on the Om10. The integration of the genetic map with the reference genome of marine medaka will serve as a good resource for studies in molecular toxicology, genomics, CRISPR/Cas9, and epigenetics.

---

Many fish species are useful for ecotoxicological research, as they indicate an early warning of environmental contamination caused by various aquatic pollutants (Arellano-Aguilar *et al.* 2009). Recently, an increase in the contamination levels in estuaries and coastal water due to anthropogenic pollutants emphasizes the needs for marine sentinel model fish species. Medaka (*Oryzias* spp.) is an important fish species in ecotoxicology and considered a model species, as its biological features include certain advantages such as small body size and short generation time (Kim *et al.* 2016). Japanese medaka *Oryzias latipes* is important and widely used model species for the studies of genetics, evolution, and ecotoxicological study, with abundant genomic resources. Since *O. latipes* is a freshwater species, their responses to environmental toxicants can be different in those of marine fish (Shi *et al.* 2008; Wu *et al.* 2012; Wheeler *et al.* 2002). Marine medaka *Oryzias melastigma* inhabits a brackish water in Asian regions including Pakistan, India, Burma, and Thailand (Naruse, 1996). *O. melastigma* has been acknowledged as a potential model fish for marine ecotoxicological studies and is useful for the evaluation of acute and/or chronic toxicity, embryo toxicity testing (Chen *et al.* 2011; Dong *et al.* 2014; Kim *et al.* 2016; Kong *et al.* 2008; Shen *et al.* 2010; Wang *et al.* 2011; Kim *et al.* 2016). Previously, *O. melastigma* was believed to be phylogenetically closely related to the Japanese medaka (*O. latipes*) (Kong *et al.* 2008; Bo *et al.* 2011; Kasahara *et al.* 2007). However, recently, *O. melastigma* and *O. latipes* were suggested to be more divergent, as they belong to two distinct species groups of medaka (Takehana *et al.* 2005). Medaka has an XX-XY sex-determining system with morphologically indistinguishable sex chromosomes (Matsuda *et al.* 1998). Phenotypic distinction between the male and female in medaka are distinguished by a number of secondary sex characters including shape and size of dorsal and anal fins (Schartl, 2004). Sex-determining gene, *dmrt1bY*, has been identified in *O. latipes*, but not in *O. celebensis* (Kondo *et al.* 2003; Matsuda *et al.* 2002). In addition, as sex-determining region Y (*Sry*) has been identified as the male-determining gene (Sinclair *et al.* 1990) and *sox3* as the ancestral precursor of *Sry* based on evolutionary evidences (Graves, 1998), it is important to understand and identify *sox3*. Furthermore, limited information on sex-determining genes in *O. melastigma* is available, thus, more genomic resources on the marine medaka (*O. melastigma*) are required.

A genetic linkage map is a very useful tool to understand genetic architecture such as chromosome structure, segregation distortion regions, recombination rate, and recombination hotspots (Myers *et al.* 2005; Zhu *et al.* 2014; Guo *et al.* 2013; Shifman *et al.* 2006). Furthermore, it provides a framework for mapping the chromosomal location of single-gene traits and quantitative traits of interest, and helps to facilitate candidate gene cloning, and comparative genomic analysis with some genome information together (Lee *et al.* 2005; Amores *et al.* 2011; Feng *et al.* 2015; Zhu *et al.* 2013; Li *et al.* 2015; Shao *et al.* 2015; Wang *et al.* 2015; Xiao *et al.* 2015; Kanamori *et al.* 2016). A high-density genetic map also plays an important role in assembling whole genome sequences by examining the accuracy of *de novo* genome assembly (Dukic *et al.* 2016; Rhee *et al.* 2017; Wang *et al.* 2017). Indeed, the importance of a high-density genetic map has been demonstrated during *de novo* genome assembly in teleost fish, as it was discovered that an additional genome duplication event had occurred (Meyer *et al.* 1999; Meyer *et al.* 2005; Volff, 2005). In addition, restriction-site associated DNA (RAD) sequencing based on next-generation sequencing (NGS) enables the rapid discovery of genome-wide genetic markers and high-throughput single nucleotide polymorphism (SNP) genotyping in mapping gene families and facilitates the construction of high-density genetic linkage maps in both model and non-model organisms (Baird *et al.* 2008; Davey and Blaxter, 2010; Amores *et al.* 2011; Davey *et al.* 2011; Etter *et al.* 2011).

We have recently published a reference genome assembly (total genome length 779.4 Mb) of *O. melastigma* as a model species in environmental toxicology (Kim *et al.* 2018). In this study, we constructed a high-density genetic linkage map of *O. melastigma* using an F1 full-sib family. Using the genetic linkage map, sex quantitative trait locus (QTL) has been mapped in the genome of *O. melastigma* and the previous genome assembly was anchored onto the linkage map to improve the contiguity of the assembly. The development of a high-density genetic map is imperative to facilitate both genetic and genomic studies in *O. melastigma.* The present study will assist in a better understanding of genome-based research in molecular toxicology, genomics, CRISPR-cas9, and epigenetics.

## MATERIALS AND METHODS

### Mapping cross

The *O. melastigma* used in this study were obtained from the City University of Hong Kong (kindly provided by Dr. Doris W.T. Au) and maintained at Nagahama Institute of Bio-Science and Technology, Nagahama, Japan. A male and a female fish were bred to produce F1 progenies. In total, 58 F1 individuals were used to create a linkage group (LG).

### RAD sequencing

Genomic DNA was extracted from muscle tissue using DNeasy Blood & Tissue Kit (Qiagen, Hilden, Germany) according to the manufacturer’s instructions. The size and quality of DNA isolated was checked on 1% agarose gene by electrophoresis and the concentration was measured using Qubit florometer (Thermo Fisher Scientific, Waltham, MA, USA). Genomic DNA (40 ng) of each sample was digested with *Bgl*II (5 Unit) and *Eco*RI (5 Unit), ligated with a Y-shaped adaptor, amplified by polymerase chain reaction (PCR) with KAPA HiFi HS ReadyMix (KAPA BIOSYSTEMS, Wilmington, MA, USA), and fragments were selected with E-Gel Size Select (Life Technologies, Carlsbad, CA, USA). The mean size of the selected fragments was 333 bp (CV 16.4%). Further details of the library preparation method are described in a previous study by Sakaguchi *et al.* (2015). RAD sequencing (RAD-seq) was performed using 58 F1 individuals and both parents with HiSeq2500 (Illumina, San Diego, CA, USA) with eight cycles for index read and 51 cycles for the reads of interest. For each parental sample, the same amounts were aliquoted in four different reaction tubes and sequencing of each reaction were carried out to reduce PCR amplification bias. All procedures related to RAD-seq and RAD-seq libraries were performed by Clockmics Inc. (Osaka, Japan).

### Extracting RAD-tags and SNP genotyping by Stacks

Quality filtration of sequence reads was performed using Trimmomatic v.0.33 (Bolger *et al.* 2014) with parameter options of −0.33.jar SE -phred33 TOPHRED33 ILLUMINACLIP TruSeq3-PE-2.fa:2:30:10 LEADING:19 TRAILING:19 SLIDINGWINDOW:30:20 AVGQUAL:20 MINLEN:51. RAD-tag extraction and genotyping were performed with Stacks v.1.47 software (http://creskolab.uoregon.edu/stacks/) (Catchen *et al.* 2011). The sequence reads were aligned to the available reference genome **(GCF_002922805.1;** Kim *et al.* 2018) using GSnap (https://www.gvst.co.uk/gsnap.htm) with default parameters (-t 30 –n 1 –m 5 –i 2), which were converted to BAM files. All RAD-tags catalog from the parental samples were extracted by Stacks using the *ref_map.pl* pipeline with the parameters –m10 and -P 3 and genotyping was called by the parameters of minimum number of 5 reads to call a homozygous genotype, a minimum minor allele frequency of 0.1 to call a heterozygote, and a maximum minor allele frequency of 0.05 to call a homozygote. Among RAD-tags, single nucleotide polymorphism (SNP) markers with maximum likelihood of 0 were selected for mapping and the SNP markers with genotypes of at least 53 F1 offsprings (> 90%) were collected for map construction using the command *genotypes* –r 53 of Stacks v.1.47. Data of raw sequences were deposited in the Sequence Read Archive (SRA) (http://www.ncbi.nlm.nih.gov/sra) under the accession number **PRJNA514812**.

### Linkage map construction

Linkage analysis of genetic markers (e.g. SNPs) was performed using JoinMap 5.0 (Wageningen, Netherlands; Van Ooijen, 2018). The SNP markers with a significant segregation distortion (χ^2^ test, *P* < 0.01) were removed from the analysis of linkage map construction. Linkage groups (LGs) were identified by the grouping parameters of independent Likelihood Odds Ratio (LOD) threshold of 5 provided in JoinMap 5.0. Map distances were calculated by the Kosambi’s mapping function and mapping algorithm used for building linkage map was regression mapping based on the default parameters. The regression mapping adds loci one by one, starting from the most informative pair of loci. The best position of each added locus is searched by comparing the goodness-of-fit of the calculated map for each tested position. JoinMap was used and performed three rounds of marker positioning with a jump threshold of 5 and we took second round of map as a final map, as recommended by the manual. The linkage map was visualized using MapCart 2.32 (Voorrips, 2002). The name of the linkage groups was matched with the homologous chromosomes of Japanese medaka.

### Anchoring the reference genome on to linkage map

Genetic markers in the linkage map were anchored to the reference genome **(GCF_002922805.1)** of the marine medaka *O. melastigma* using Chromonomer v. 1.08 (http://catchenlab.life.illinois.edu/chromonomer/). The integrated genome assembly based on the genetic map was re-assessed with benchmarking universal single-copy orthologs (BUSCO) v.3.0 (Simao *et al.* 2015) using the vertebrate database (OrthoDB v.9.0; https://www.orthodb.org/?page=filelist). The gene annotation of the final assembly using MAKER v.2.31.8 pipeline with manual curation (**Suppl. Fig. S1)** (Holt and Yandell, 2011).

### Comparative analysis of two medaka genomes

The final linkage map based on the genome assembly of marine medaka was compared with the genome of Japanese medaka (Hd-rR strain) to compare the similarity between two medaka genomes using Mummer v.3.0 (http://mummer.sourceforge.net/manual/#coords).

### Sex linkage analysis

The mapping panel consisted of 27 males and 31 females. Since the sex trait is qualitative, the phenotype was converted into binary code; 1 indicating male and 0 indicating female. To analyze sex linkage, standard interval mapping was performed and significance was determined by permutation test (n=1000) using MapQTL 6.0 (van Ooijen *et al.* 2002).

### Data availability

Sequenced species *O. melastigma* is available upon request. Suppl. Data contains all supplemental data and figures. **Suppl. File S1** contains the names of 2481 SNP, genotype, map positions, and tag sequences in Map info tab and phenotypes for sex of all individuals were in Phenotype tab. **Suppl. File S2** contains the re-scaffolding process and results by Chromonomer. Raw data of RAD seq were deposited to GenBank under the accession number **PRJNA514812**. The genome sequence data are available in GenBank with accession number **PRJNA401159**. The genome JBrowse is available at http://rotifer.skku.edu:8080/Om1. Supplemental materials are available at **Figshare**.

## RESULTS

### Constructing genetic linkage map of marine medaka *Oryzias melastigma*

Stacks software extracted 113,367 RAD-tags from *O. melastigma* gnome. Among them, the number of putative SNP markers was 34,040. To distinguish polymorphisms and sequencing errors, we collected 24,441 SNP markers with Likelihood Ratio of 0, including 8,518 SNP markers that have been located in the same RAD-tags more than twice. Among the remaining 15,922 loci, 4,497 SNP loci were successfully genotyped in at least 53 F1 individuals (>90%). After removing the markers that showed a segregation distortion (*P*<0.01), 3,732 were finally used for building a genetic map. The 3,730 markers were grouped into 24 LGs with a LOD ≥ 5.0 and two remaining markers were not linked with any of those groups. The regression mapping function by JoinMap positioned 2,481 SNPs in the second rounds of mapping by comparing the goodness-of-fit (**Suppl. File S1**). The 24 LGs were consistent with the number of chromosome (n=24) in *O. melastigma* (Uwa *et al.* 1983). The total map length was 1,784 cM and each LG included 57-173 markers with an average marker interval of 0.72 cM (**Figure 1** and **Table 1**).

**Table 1.** Summary of the genetic linkage map of *Oryzias melastigma*.

| LG | Group ID | No. of markers<br>mapped | Map Length (cM) | *Oryzias latipes*<br>chromosome |
| --- | --- | --- | --- | --- |
| Om01 | Group 7 | 91 | 77.92 | 1 |
| Om02 | Group 21 | 98 | 64.935 | 2 |
| Om03 | Group 5 | 107 | 89.184 | 3 |
| Om04 | Group 6 | 108 | 68.313 | 4 |
| Om05 | Group 13 | 102 | 84.132 | 5 |
| Om06 | Group 4 | 173 | 89.972 | 6 |
| Om07 | Group 12 | 118 | 80.081 | 7 |
| Om08 | Group 24 | 57 | 53.584 | 8 |
| Om09 | Group 17 | 111 | 85.671 | 9 |
| Om10 | Group 16 | 117 | 68.297 | 10 |
| Om11 | Group 1 | 103 | 79.839 | 11 |
| Om12 | Group 8 | 123 | 78.25 | 12 |
| Om13 | Group 15 | 74 | 71.657 | 13 |
| Om14 | Group 14 | 142 | 81.03 | 14 |
| Om15 | Group 18 | 106 | 57.118 | 15 |
| Om16 | Group 19 | 88 | 53.324 | 16 |
| Om17 | Group 3 | 128 | 77.589 | 17 |
| Om18 | Group 2 | 117 | 72.043 | 18 |
| Om19 | Group 22 | 99 | 72.178 | 19 |
| Om20 | Group 23 | 83 | 93.802 | 20 |
| Om21 | Group 20 | 93 | 66.785 | 21 |
| Om22 | Group 9 | 79 | 72.197 | 22 |
| Om23 | Group 11 | 71 | 72.7 | 23 |
| Om24 | Group 10 | 93 | 73.366 | 24 |
|  |  | 2,481 | 1783.967 |  |

**Figure 1.**
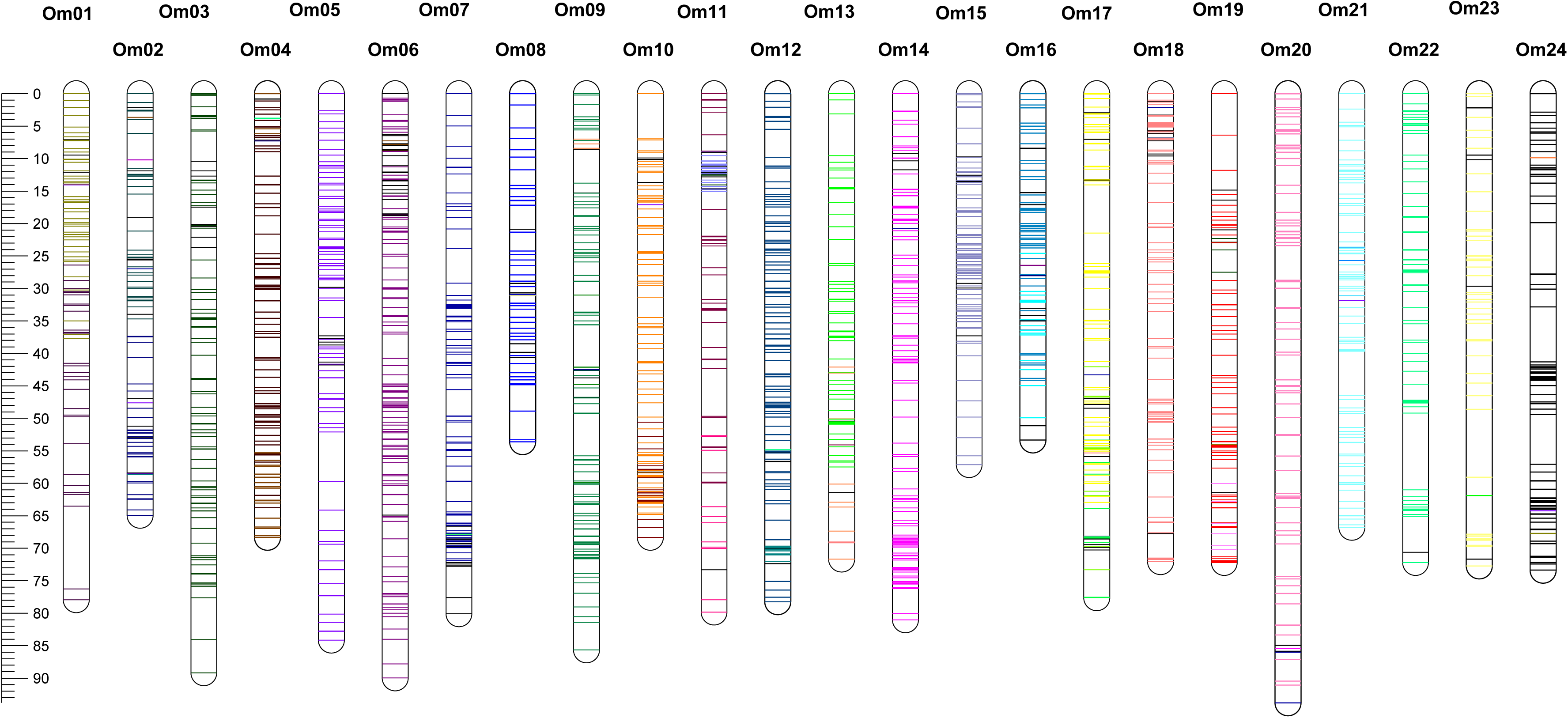
A linkage map of the marine medaka *Oryzias melastigma.* The map consists of 24 linkage groups and the bars on each linkage group single nucleotide polymorphism (SNP) markers. Colors of bars indicate the reference-assisted chromosome assembly scaffolds that SNP was extracted. Name, sequences, and position of SNP are included in the **Suppl. File 1**.

### Re-scaffolding of the reference genome with the genetic map

Using the SNP marker information in the high-density genetic linkage map, 810 markers were anchored to 134 scaffolds and among them, 35 were split into 1 to 9 positions, producing a total of 260 integrated scaffolds (**Tables 2, Suppl. Table** S**2**, and **Suppl. Fig. S2**). After integration, the length of the genome scaffolds aligned on the map was 712,537,413 bp (**Table 2**). Out of the 260 integrated scaffolds, the orientation was determined in 160 scaffolds spanning 670,530,120 bp, which accounted for 94% of the total scaffold length in the linkage map (**Table 2**). Among 40 reference-assisted chromosome assembly (RACA) scaffolds previously published (Kim *et al.* 2018), 20 RACA scaffolds (RACA3, 5, 6, 7, 8, 9, 11, 12, 14, 15, 18, 21, 22, 23, 25, 26, 27, 29, 30, 40) were aligned to the 13 linkage groups (Om23, Om21, Om20, Om19, Om17, Om16, Om14, Om 12, Om11, Om10, Om09, Om08, Om01) without any modification (**Suppl. Table S1**). Other RACA scaffolds showed the major alignment of one linkage group with the partial alignment of another linkage groups (**Suppl. Table S1**). Among them, RACA 31 showed the most frequent rearrangement during the anchoring process, which was largely aligned with Om 7 and partially aligned with another 4 linkage groups (Om20, Om18, Om17, and Om02) (**Suppl. Table S1**). Four linkage groups (Om03, Om05, Om15, and Om22) were completely aligned by each of four RACA scaffolds (RACA4, RACA16, RACA33, and RACA36), respectively (**Table 2**), although some parts of the aligned sequences in the linkage groups were inserted from small scaffolds (**Suppl. File S2**) and the parts of RACA scaffolds were located in parts of another linkage groups (**Supp. Table S1**). Overall, the final genetic map based genome assembly consisted of 8,493 scaffolds; 24 linkage map-based scaffolds (90.7%) and 8,469 unanchored scaffolds (9.3%). The total genome length was 778.7 Mb with an N50 value of 29,978,720 bp (**Table 3**). BUSCO analysis indicated that the final genome assembly of *O. melastigma* represented 96.8% of the complete copy in the vertebrate gene model (**Table 4**). The genome annotation pipeline in the final assembly was defined as 24,506 genes (http://rotifer.skku.edu:8080/Om1), ranging from 661 to 1,216 genes per LG (**Table 3** and **Suppl. Table S3**).

**Table 2.**
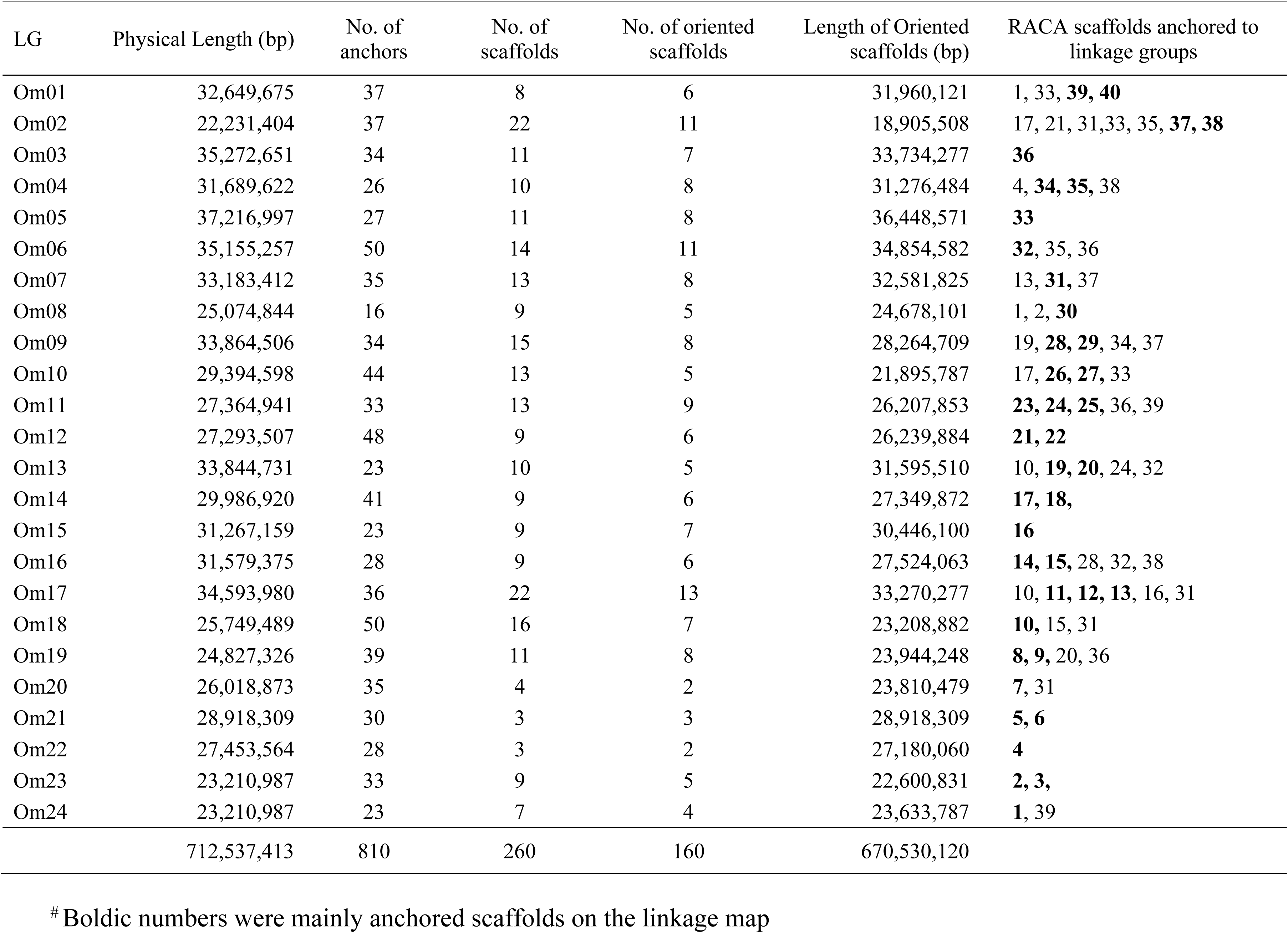
Physical lengths of linkage map anchored with the reference genome assembly in *Oryzias melastigma*^*#*^.

| LG | Physical Length (bp) | No. of anchors | No. of scaffolds | No. of oriented scaffolds | Length of Oriented scaffolds (bp) | RACA scaffolds anchored to linkage groups |
| --- | --- | --- | --- | --- | --- | --- |
| Om01 | 32,649,675 | 37 | 8 | 6 | 31,960,121 | 1, 33, **39, 40** |
| Om02 | 22,231,404 | 37 | 22 | 11 | 18,905,508 | 17, 21, 31,33, 35, **37, 38** |
| Om03 | 35,272,651 | 34 | 11 | 7 | 33,734,277 | **36** |
| Om04 | 31,689,622 | 26 | 10 | 8 | 31,276,484 | 4, **34, 35**, 38 |
| Om05 | 37,216,997 | 27 | 11 | 8 | 36,448,571 | **33** |
| Om06 | 35,155,257 | 50 | 14 | 11 | 34,854,582 | **32**, 35, 36 |
| Om07 | 33,183,412 | 35 | 13 | 8 | 32,581,825 | 13, **31**, 37 |
| Om08 | 25,074,844 | 16 | 9 | 5 | 24,678,101 | 1, 2, **30** |
| Om09 | 33,864,506 | 34 | 15 | 8 | 28,264,709 | 19, **28, 29**, 34, 37 |
| Om10 | 29,394,598 | 44 | 13 | 5 | 21,895,787 | 17, **26, 27**, 33 |
| Om11 | 27,364,941 | 33 | 13 | 9 | 26,207,853 | **23, 24, 25**, 36, 39 |
| Om12 | 27,293,507 | 48 | 9 | 6 | 26,239,884 | **21, 22** |
| Om13 | 33,844,731 | 23 | 10 | 5 | 31,595,510 | 10, **19, 20**, 24, 32 |
| Om14 | 29,986,920 | 41 | 9 | 6 | 27,349,872 | **17, 18**, |
| Om15 | 31,267,159 | 23 | 9 | 7 | 30,446,100 | **16** |
| Om16 | 31,579,375 | 28 | 9 | 6 | 27,524,063 | **14, 15**, 28, 32, 38 |
| Om17 | 34,593,980 | 36 | 22 | 13 | 33,270,277 | 10, **11, 12, 13**, 16, 31 |
| Om18 | 25,749,489 | 50 | 16 | 7 | 23,208,882 | **10**, 15, 31 |
| Om19 | 24,827,326 | 39 | 11 | 8 | 23,944,248 | **8, 9**, 20, 36 |
| Om20 | 26,018,873 | 35 | 4 | 2 | 23,810,479 | **7**, 31 |
| Om21 | 28,918,309 | 30 | 3 | 3 | 28,918,309 | **5, 6** |
| Om22 | 27,453,564 | 28 | 3 | 2 | 27,180,060 | **4** |
| Om23 | 23,210,987 | 33 | 9 | 5 | 22,600,831 | **2, 3**, |
| Om24 | 23,210,987 | 23 | 7 | 4 | 23,633,787 | **1**, 39 |
|  | 712,537,413 | 810 | 260 | 160 | 670,530,120 |  |

<sup>#</sup> Boldic numbers were mainly anchored scaffolds on the linkage map

**Table 3.** Statistics of the final genome assembly before and after anchoring in *Oryzias melastigma*.

| Statistics | Reference genomes |  |
| --- | --- | --- |
|  | RACA | Linkage map based |
| Number of scaffolds<sup>#</sup> | 8,602 | 8,493 |
| Length of scaffolds (bp) | 779,456,607 | 778,703,520 |
| N50 (bp) | 23,737,187 | 29,987,720 |
| Largest scaffold (bp) | 37,948,421 | 37,217,997 |
| Gap (%) | 5.29 | 5.2 |
| GC content (%) | 39.04 | 37.02 |
| Number of unanchored scaffolds |  | 8,469 |
| Length of unanchored scaffolds (bp) |  | 66,142,507 |
| Number of genes | 23,528 | 24,506 |
| Total genes length (bp) | 51,834,196 | 24,784,506 |
| Average genes length (bp) | 2,586 | 1,011 |
| Maximum gene length (bp) | 80,775 | 26,364 |
| GC content (%) | 54.34 | 54.27 |

**Table 4.** Assessment of LG-based assembly completeness.

| Vertebrata database | % | Numbers of genes<br>(n=2586) |
| --- | --- | --- |
| Complete BUSCOs (C) | 96.8 | 2504 |
| Complete and single-copy BUSCOs (S) | 95.8 | 2477 |
| Complete and duplicated BUSCOs (D) | 1 | 27 |
| Fragmented BUSCOs (F) | 1.2 | 32 |
| Missing BUSCOs (M) | 2 | 50 |

### Comparative genomic analysis of two medaka genomes

The final genome assembly, with an integration of a genetic map, provides an efficient resource for comparative genomic analysis with other medaka genomes such as a Japanese medaka (*O. latipes*). The 24 genetic map-based scaffolds showed good homology in gene contents and sequences similarity with chromosomes in *O. latipes*. Most LG-based scaffolds of marine medaka showed collinear relationships completely or in a majority with the counterpart chromosomes of *O. latipes* **(Fig. 2** and **Suppl. Fig. S3).** Other LG/chromosomes in *O. melastigma* showed disrupted collinearity due to intrachromosomal rearrangements or possible errors in linkage map. Reversely matched parts in the collinearly related scaffolds were most likely caused by the undetermined orientation of scaffolds during the anchoring process. In addition, some LG-based scaffolds showed in/del regions (Om05, Om08, Om11, Om14, Om15, Om16, and Om17) compared to Japanese medaka chromosomes.

**Figure 2.**
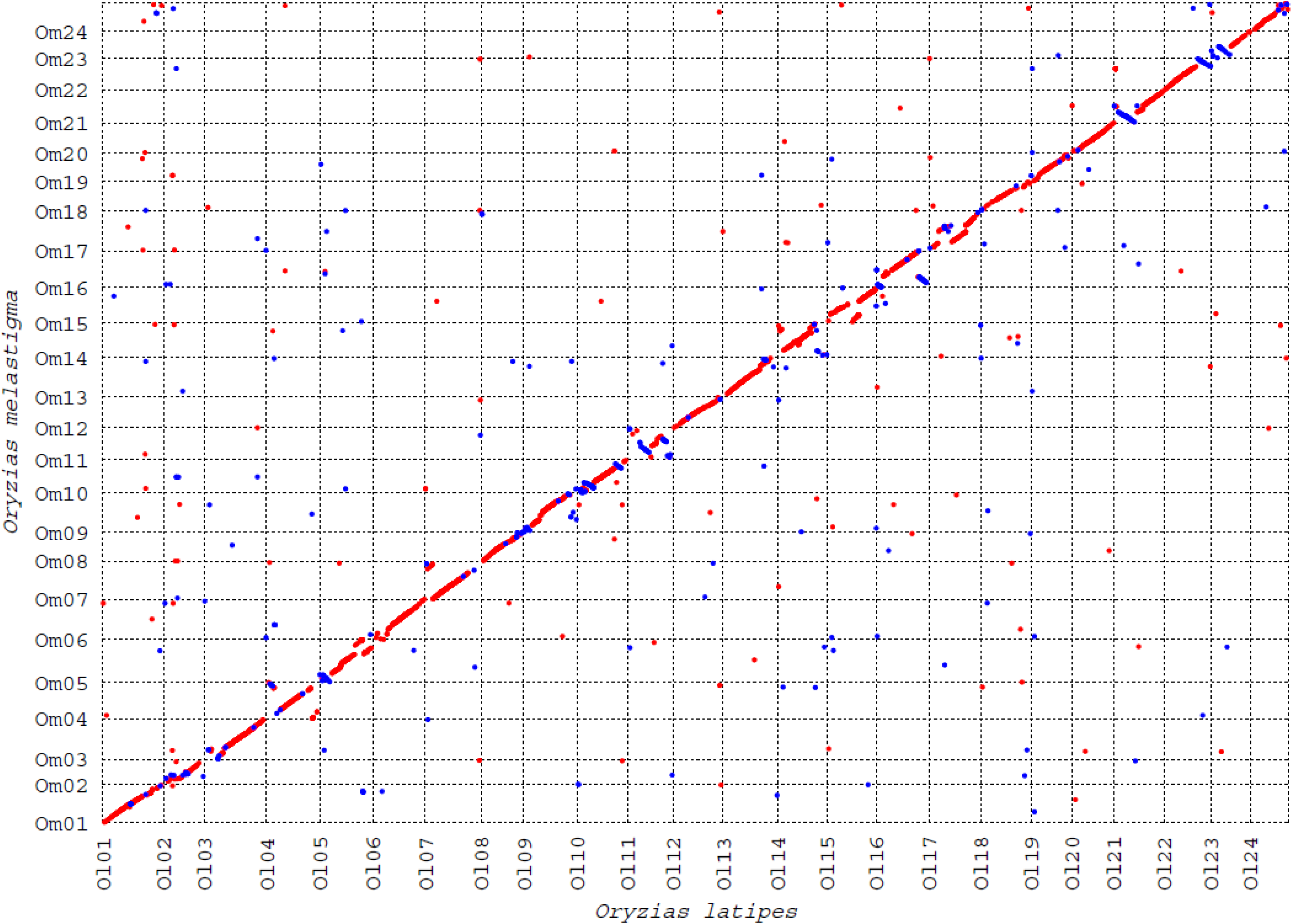
Genome-wide comparison of the genomic position of homologous loci between *Oryzias melastigma* and *O. latipes* using Mummer. Red and blue dots represent forward and reverse match, respectively.

### Mapping of sex-determining regions

A significant QTL for sex was detected in Om10 with the LOD significance threshold of 5.3 based on the permutation test (**Fig. 3**). Among 2,481 markers mapped in the Om linkage groups, 58 markers showed the significant association with sex (**Suppl. Table S4**), most of which were markers from the RACA27. The two most significant markers associated with sex were 17040 (24.3 cM) and 17018 (24.4 cM) with LOD of 35. The genotypes of those markers were completely linked with the sex phenotype of all individuals, except for one animal (d004), which could be caused by wrong phenotyping or sex-reversal. Gene contents in this region, however, showed no obvious candidate gene for sex determination.

**Figure 3.**
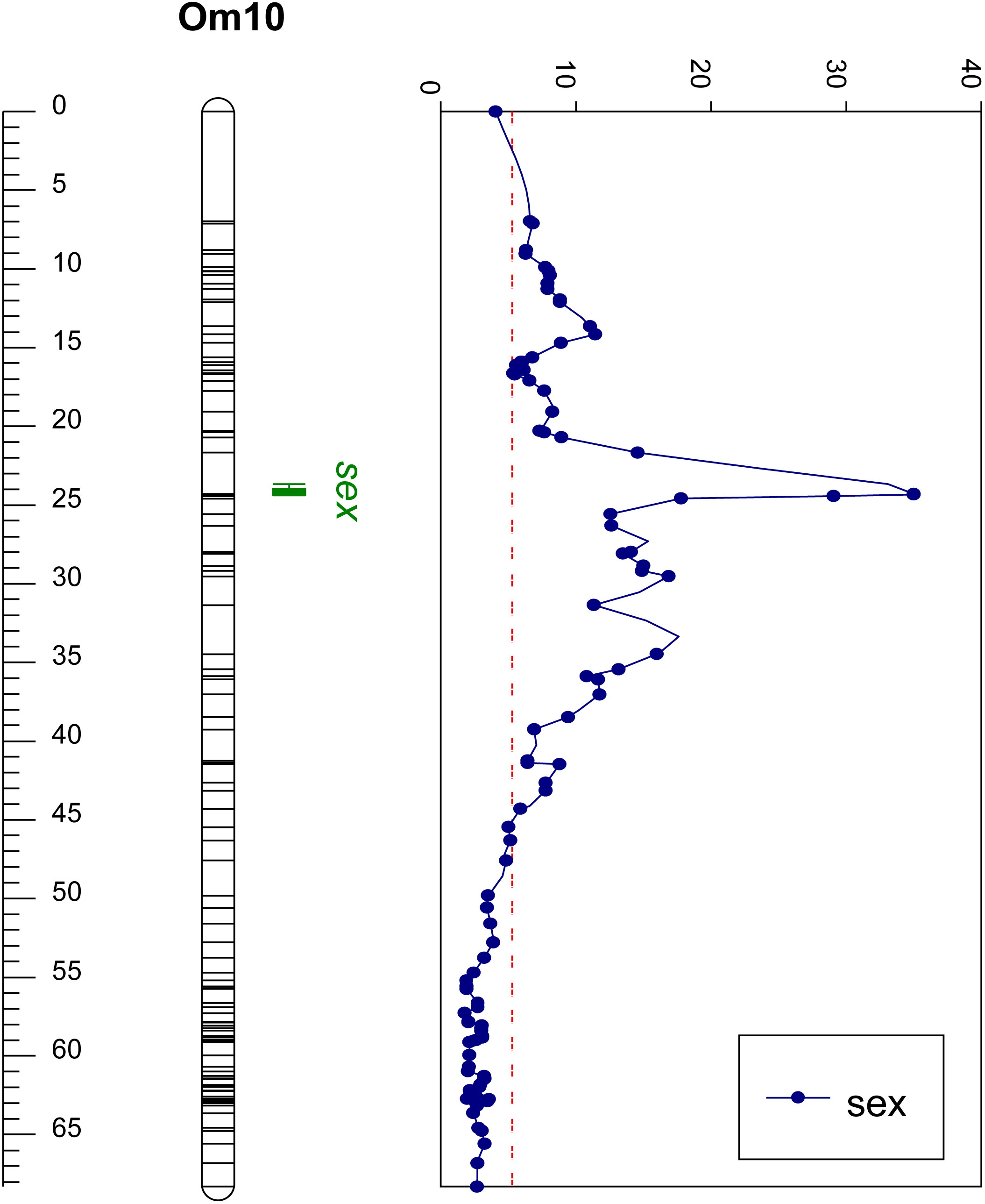
Quantitative trait locus mapping for sex traits on the Om10 scaffold of *Oryzias melastigma*. Scales on the left represents the map position (cM) of Om10, whereas scales on the top of the graph represent the value of the likelihood odds ratio (LOD) scores. The red dotted line indicates the threshold of significance (LOD=5.3).

## DISCUSSION

A high-resolution genetic map is very useful in diverse genomic research and has been applied to many fish species (Amores *et al.* 2011; Li *et al.* 2015; Shao *et al.* 2015; Xiao *et al.* 2015; Wang *et al.* 2015; Kanamori *et al.* 2016). In this study, a high-density genetic map of *O. melastigma* was constructed using RAD sequencing and was used for verifying the previously published marine medaka reference genome and for aligning the scaffolds at the chromosomal level. The genetic map of *O. melastigma* consists of 24 LGs with 2,481 SNP markers, which cover 24 chromosomes (Uwa *et al.* 1983). The 810 genetic markers were anchored to the genetic map (**Table 2** and **Suppl. Fig. 2**), which mapped 90.7% (713 Mb) of the reference genome assembly sequence onto the genetic map (**Table 3**). Although all SNP markers were extracted by the alignment of RAD sequences against the reference genome of marine medaka (Kim *et al.* 2018), the number of anchored markers were lower than anticipated. Many markers in the LGs were excluded from the integrating process due to inconsistency between the genetic map and reference assembly, indicating possible errors in marker order and/or *de novo* assembly. Indeed, in most cases, the markers were tended to be highly concentrated in the narrow regions, suggesting that more recombinant individuals will be required to obtain a more precise marker order. It is likely that the size of the mapping family was not big enough for the marker concentrated regions, compared with the number of markers analyzed. Despite this shortage, the marine medaka genetic map was still integrated with the reference assembly, considering other fish species had a similar or slightly lower level of scaffolds mapping onto the genetic map (Tine *et al.* 2014; Li *et al.* 2015; Wang *et al.* 2017).

Previously, we have developed a reference genome for the marine medaka *O. melastigma* in several steps (Kim *et al.* 2018). *De novo* assembly was performed using the combination of Platanus and Haplomerger2 assemblers (Kajitani *et al.* 2014; Huang *et al.* 2017). The contiguity of *de novo* genome assembly was further increased by RACA (Kim *et al.* 2013), which assisted in the construction of highly ordered and oriented scaffolds of *de novo* assembly and reassembled scaffolds into longer chromosomal fragments using the comparative genome information of closely related species. RACA assembly of *O. melastigma* generated 40 gigantic scaffolds (**Suppl. Table 1**), which account for 674 Mb (86.5%) of the total genome sequence (Kim *et al.* 2018). Taken together, the genetic map-integrated assembly improved the reference sequences by mapping 90.7% of genome sequences onto 24 LGs with 94% determined orientation of mapped scaffolds **(Table 2)**. The BUSCO and N50 demonstrated a better quality of integrated final genome assembly with values increased to 96.8% and 29,987,720 bp, respectively **(Tables 3 and 4)**.

Quantitative trait locus (QTL) analysis identified that Om10 was strongly associated with sex in *O. melastigma*. Since comparative syntenic analysis showed that this homologous region showed a collinear relationship between *O. latipes* and *O. melastigma* (**Fig. 2**) but no obvious candidate genes were in the sex QTL region, we investigated the gene list in the corresponding region of *O. latipes* to examine missing genes during the gene annotation processes in marine medaka. Based on the gene contents in *O. latipes, sox3* was located in the homologous region but in *O. melastigma, sox3* was located in Om14 (**Suppl. Fig. 4**). We took a closer look at this region in *O. melastigma* to determine if any *sox*-like genes were located in this homologous region but the existence of the undetemined Ns indicated that there were gaps in the sequence, which were possibly caused by repeats. It was unlikely that Ns were produced by the Chromonomer, as those whole sex-determining regions were on the RACA scaffold 27. It is well known that repetitive DNA accumulates in sex-determining regions in fish (Oliveira *et al.* 1999; Harvey *et al.* 2002; Matsuda *et al.* 2002). For further analysis, we need to narrow down the QTL regions with fine mapping and the functional analysis of candidate genes in the region.

In summary, a high-density genetic map is a very useful resource to determine the accuracy of *de novo* genome assembly, especially with massively parallel short sequencing reads. In addition, the integration of a genetic map with reference genomes will be a useful resource in various genomic studies including comparative genomic analyses, fine mapping of QTL, positional cloning of candidate genes, and CRISPR/Cas9 studies.

## ACKNOWLEDGEMENTS

We thank two anonymous reviewers for their valuable comments to improve the manuscript. This study was supported by a grant from the National Research Foundation (2017R1D1A1B03036026) to Bo-Young Lee.

